# Tolerance to hypothermic and antinoceptive effects of Δ9-tetrahydrocannabinol (THC) vapor inhalation in rats

**DOI:** 10.1101/172759

**Authors:** Jacques D. Nguyen, Yanabel Grant, Tony M. Kerr, Arnold Gutierrez, Maury Cole, Michael A. Taffe

**Author notes:** Address Correspondence to: Dr. Michael A. Taffe, Department of Neuroscience, SP30-2400; 10550 North Torrey Pines Road; The Scripps Research Institute, La Jolla, CA 92037; USA.

## Abstract

**Rationale:** A reduced effect of a given dose of Δ^9^-tetrahydrocannabinol (THC) emerges with repeated exposure to the drug. This tolerance can vary depending on THC dose, exposure chronicity and the behavioral or physiological measure of interest. A novel THC inhalation system based on e-cigarette technology has been recently shown to produce the hypothermic and antinociceptive effects of THC in rats.

**Objective:** To determine if tolerance to these effects can be produced with repeated vapor inhalation.

**Methods:** Groups of male and female Wistar rats were exposed to 30 minutes of inhalation of the propylene glycol (PG) vehicle or THC (200 mg/mL in PG) two or three times per day for four days. Rectal temperature changes and nociception were assessed after the first exposure on the first and fourth days of repeated inhalation.

**Results:** Female, but not male, rats developed tolerance to the hypothermic and antinociceptive effects of THC after four days of twice-daily THC vapor inhalation. Thrice daily inhalation for four days resulted in tolerance in both male and female rats. The plasma THC levels reached after a 30 minute inhalation session did not differ between the male and female rats.

**Conclusions:** Repeated daily THC inhalation induces tolerance in female and male rats, providing further validation of the vapor inhalation method for preclinical studies.

**Abbreviations:** PG, propylene glycol; THC; Δ^9^tetrahydrocannabinol;

## 1. Introduction

Human cannabis consumers are increasingly using e-cigarette type devices filled with cannabis extracts to administer an active dose of Δ^9^-tetrahydrocannabinol (THC) and other constuents (Mammen et al., 2016; Morean et al., 2015; Morean et al., 2017). This has spurred interest in the development of pre-clinical models which are capable of a similar route of drug administration in laboratory rodents. Recent studies showed that intrapulmonary delivery of THC using an e-cigarette based system results in a robust and dose-dependent hypothermia and antinociception in male and female rats (Nguyen et al., 2016b) and a similar approach produces hypothermia in mice following inhalation exposure to synthetic cannabinoid agonists (Lefever et al., 2017). Such systems also induce locomotor stimulation after inhalation of methamphetamine, mephedrone or 3,4-methylenedioxypyrovalerone (Nguyen et al., 2016a) and can produce escalated self-administration of sufentanil to the point of dependence (Vendruscolo et al., 2017). These prior studies of cannabinoid inhalation focused on the acute effects of THC and did not attempt to provide any evidence of plasticity (i.e., tolerance or sensitization) of the cannabinoid-typical behavioral or physiological effects that result. It has been well established that repeated exposure to THC via parenteral injection produces tolerance to the acute effects in laboratory animals such as mice (Anderson et al., 1975; Fan et al., 1994), rats (Jarbe, 1978; Taylor and Fennessy, 1978; Wiley and Burston, 2014) or monkeys (Ginsburg et al., 2014; Smith et al., 1983; Winsauer et al., 2011). Thus it is of significant interest to determine if repeated exposure to THC via e-cigarette type inhalation is capable of producing tolerance in rats to further validate this model.

A substantial and sustained decrease in body temperature, and a reduction in sensitivity to a noxious stimulus (antinociception), are two major indicators of cannabinoid-like activity in laboratory rodents (Wiley et al., 2014). These measures have been shown to be consistently affected by the inhalation of THC via e-cigarette type technology in both male and female rats (Javadi-Paydar et al., 2017; Nguyen et al., 2016b). This study was therefore designed to determine if the body temperature response to, and antinociceptive effects of, the inhalation of THC via e-cigarette type technology are attenuated by repeated daily exposure in male or female rats. As humans increasingly ingest THC via e-cigarette technology it has become increasingly important to develop translational pre-clinical models that are capable of evaluating the consequences of this route of administration. This study of THC-induced tolerance to the effects of THC inhalation is therefore a critical advance.

## 2. Methods

### 2.1. Subjects

Male (N=16) and female (N=16) Wistar (Charles River, New York) rats were housed in humidity and temperature-controlled (23±2 °C) vivaria on 12:12 hour light:dark cycles. Rats had *ad libitum* access to food and water in their home cages and all experiments were performed in the rats’ scotophase. Rats entered the laboratory at 10-11 weeks of age. All procedures were conducted under protocols approved by the Institutional Animal Care and Use Committee of The Scripps Research Institute.

### 2.2 Drugs

Δ^9^-tetrahydrocannabinol (THC) was administered by vapor inhalation with doses described by the concentration in the propylene glycol (PG) vehicle and duration of inhalation. The THC was provided by the U.S. National Institute on Drug Abuse.

### 2.3 Inhalation Apparatus

Sealed exposure chambers for the first study were modified from a 254mm × 267mm × 381mm Allentown, Inc (Allentown, NJ) rat cage to regulate airflow and the delivery of vaporized drug to rats as has been previously described (Nguyen et al, 2016a; Nguyen et al, 2016b). An e-vape controller (Model SSV-1; La Jolla Alcohol Research, Inc, La Jolla, CA, USA) was triggered to deliver the scheduled series of puffs from Protank 3 Atomizer (Kanger Tech; Shenzhen Kanger Technology Co.,LTD; Fuyong Town, Shenzhen, China) e-cigarette cartridges by MedPC IV software (Med Associates, St. Albans, VT USA).

Sealed exposure chambers (152mm × 178 mm × 330 mm) for the second study were manufactured by LJARI. An e-vape controller (Model SSV-3; 58.0 watts; La Jolla Alcohol Research, Inc, La Jolla, CA, USA) was scheduled to deliver 6 second puffs every five minutes from a Smok Baby Beast Brother TFV8 sub-ohm tank (fitted with the V8 X-Baby M2 0.25 ohm coil). The parameters for the second generation apparatus were selected to produce similar acute effects of THC inhlation as the first study based on pilot and other ongoing studies.

The chamber air for both apparatuses was vacuum-controlled by a chamber exhaust valve (i.e., a “pull” system) to flow room ambient air through an intake valve at ∼2-3 L per minute. This functioned to ensure that vapor entered the chamber on each device-triggered event. The vapor stream was integrated with the ambient air stream once triggered. The vapor cloud cleared from the chamber by 4 minutes after the puff delivery.

### 2.4 Body Temperature

The body temperature was determined by rectal measurement with a lubricated thermistor (VWR Traceable^TM^ Digital Thermometer) as previously described (Gilpin et al., 2011).

### 2.5 Nociception Assay

Tail-withdrawal antinociception was assessed using a water bath (Bransonic® CPXH Ultrasonic Baths, Danbury, CT) maintained at 52°C. The latency to withdraw the tail was measured using a stopwatch, and a cutoff of 15 seconds was used to avoid any possible tissue damage (Wakley and Craft, 2011; Wakley et al, 2014). Tail-withdrawal was assessed 35, 60 and 120 minutes after the initiation of inhalation for the first study and 35 minutes after the start of inhalation for the second study. The person performing the assay was blinded to the treatment condition for a given subject.

### 2.6. Plasma THC Analysis

Blood samples were collected (∼500 ul) via jugular needle insertion under anesthesia with an isoflurane/oxygen vapor mixture (isoflurane 5% induction, 1–3% maintenance) 35 minutes post-initiation of vapor inhalation. Plasma THC content was quantified using fast liquid chromatography/mass spectrometry (LC/MS) adapted from (Irimia et al., 2015; Lacroix and Saussereau, 2012; Nguyen et al., 2017). 5 μL of plasma were mixed with 50 μL of deuterated internal standard (100 ng/mL CBD-d3 and THC-d3; Cerilliant), and cannabinoids were extracted into 300 μL acetonitrile and 600 μL of chloroform and then dried. Samples were reconstituted in 100 μL of an acetonitrile/methanol/water (2:1:1) mixture. Separation was performed on an Agilent LC1100 using an Eclipse XDB-C18 column (3.5um, 2.1mm × 100mm) using gradient elution with water and methanol, both with 0.2 % formic acid (300 μL/min; 73-90%). Cannabinoids were quantified using an Agilent MSD6140 single quadrupole using electrospray ionization and selected ion monitoring [CBD (m/z=315.2), CBD-d3 (m/z=318.3), THC (m/z=315.2) and THC-d3 (m/z=318.3)]. Calibration curves were conducted for each assay at a concentration range of 0-200 ng/mL and observed correlation coefficients were 0.999.

### 2.6 Data Analysis

Data (rectal temperature, tail-withdrawal latency) were analyzed with two-way Analysis of Variance (ANOVA) including repeated measures factors for the Drug treatment and Time Post-Initiation (Experiment 1 only) and a between-groups factor of Sex. Any significant main effects were followed with post-hoc analysis using the Tukey (multi-level comparisons) or Sidak (two-level comparisons) procedures. All analysis used Prism 7 for Windows (v. 7.03; GraphPad Software, Inc, San Diego CA).

### 2.7 Experiments

#### 2.7.1 Experiment 1 (Twice-daily THC vapor inhalation)

The initial study was conducted in groups of male (N=8; 17 weeks of age at start of this study) and female (N=8; 17 weeks of age) Wistar rats. Inhalation sessions were 30 minutes in duration with four 10 second vapor puffs delivered every 5 minutes with a 5.25 h interval between session initiations on each repeated-exposure day. The THC concentration was 200 mg/ml in this study. These animals had been initially evaluated with parenteral injection of THC (0, 10, 20, 20 mg/kg i.p.) in the stated (ascending) order with 2-3 day intervals between test days completed six weeks prior to the start of the chronic vapor inhalation experiment (not shown). For the repeated dosing, rats were exposed to vapor inhalation for 4 sequential days with twice-daily sessions (4 h interval between session initiations each day). Animals were evaluated for temperature and nociceptive responses after one PG-inhalation session ∼4 weeks after repeated dosing. Temperature and antinociceptive effects of THC inhalation were assessed starting ∼35 min after the start of inhalation on the first and seventh sessions.

#### 2.7.2 Experiment 2 (Thrice-daily THC vapor inhalation)

The second study was conducted with groups of experimentally naive female and male rats (N=8 per group) aged 14 weeks at the start of the experiments. The main goal was to use a more intensive THC regimen to determine if that was sufficient to induce tolerance in the males, after the failure to observe tolerance in Experiment 1. Rats first received a baseline temperature / nociception assessment without any inhalation treatment. Next, rats were assessed after PG inhalation and then after the 1^st^ and 10^th^ exposures during the THC regimen of four days with thrice-daily inhalation (3 h interval between session initiations on each day). Inhalation sessions were 30 minutes in duration with 1 six second vapor puff of THC (200 mg/ml) delivered every 5 minutes.

#### 2.7.3 Experiment 3 (Plasma THC levels)

This study was conducted in the second groups of male and female rats that were exposed to repeated THC three times-daily sessions in Experiment 2. Blood was withdrawn ∼35 min after the start of inhalation of THC (200 mg/mL). Blood collections were conducted seven days (N= 4 per group) or 17 days (N=4 per group) after the last session of the repeated-dosing experiment.

## 3. Results

### 3.1 Experiment 1 (Twice-daily THC vapor inhalation)

The inhalation of THC (200 mg/mL in the PG; 30 minute exposure) reduced body temperature and increased tail-withdrawal latency in male and female rats (Figure 1). Statistical analysis of female rectal temperature confirmed significant effects of Time [F (3, 21) = 19.85; P<0.0001], Drug treatment condition [F (2, 14) = 58.15; P<0.0001] and the interaction of factors [F (6, 42) = 14.08; P<0.0001]. The Tukey post-hoc test confirmed that rectal temperature was significantly lower after the first THC session compared with the PG session (30-120 minutes post-initiation) or with the seventh THC session (30-60 minutes post-initiation), and lower after the seventh THC session relative to vehicle (30 minutes). Statistical analysis of the male rectal temperature similarly confirmed main effects of Time [F (3, 21) = 6.96; P<0.005], Drug treatment condition [F (2, 14) = 20.91; P<0.0001] and the interaction of factors [F (6, 42) = 4.69; P<0.005]. In this case the Tukey post-hoc analysis confirmed that rectal temperature was significantly lower after the first (30-60 minutes post-initiation) and seventh (30-240 minutes post-initiation) THC sessions compared with the PG session but did not significantly differ between the two THC sessions.

**Figure 1:**
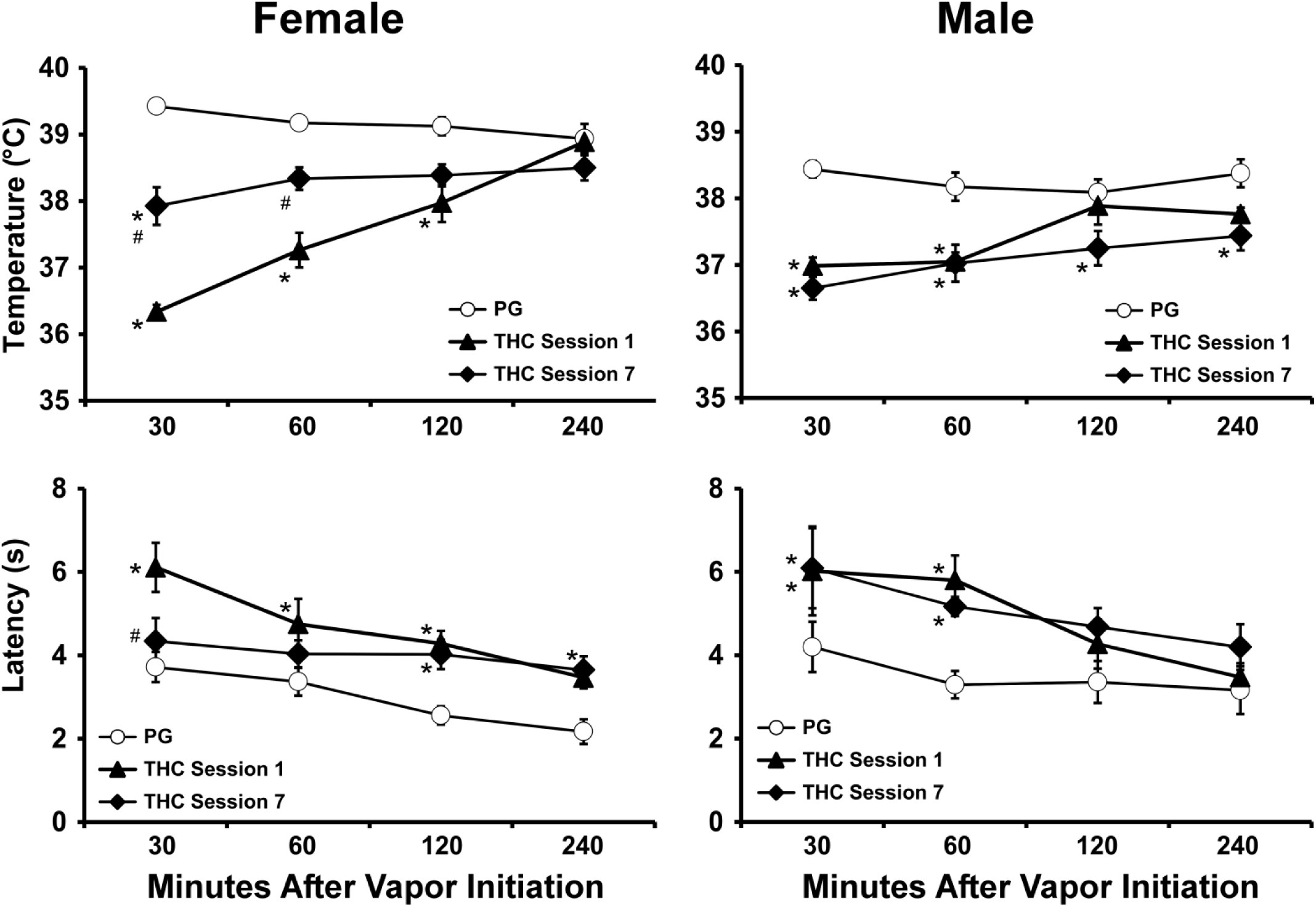
Mean (N=8 per group; ±SEM) rectal temperature (upper panels) and tail-withdrawal latency (lower panels) for male and female rats after inhalation of the PG vehicle or THC (200 mg/mL) on the 1^st^ or 7^th^ repeated session. Significant differences from the PG condition are indicated with * and from the first THC Session with #.

THC inhalation also increased tail-withdrawal latencies in both male and female groups (Figure 1). The analysis of the female withdrawal latencies confirmed significant effects of Time [F (3, 21) = 10.79; P<0.0005], Drug treatment condition [F (2, 14) = 10.5; P<0.005] and the interaction of factors [F (6, 42) = 2.51; P<0.05]. The Tukey post-hoc confirmed that withdrawal latency was longer in the first THC session compared with the PG session (30-120 minutes post-initiation) or the seventh THC session (30 minutes post-initiation) and shorter after PG inhalation relative to the seventh THC session (120-240 minutes). Analysis of the male rat tail-withdrawal latency confirmed main effects of time [F (3, 21) = 6.13; P<0.005] and drug treatment condition [F (2, 14) = 4.6; P<0.05] but not of the interaction. The Tukey post-hoc on the marginal means confirmed that withdrawal latency was significantly longer in the seventh THC session compared with vehicle but did not differ between the first and seventh THC sessions.

An analysis of the tail-withdrawal latencies 30 minutes after initiation of inhalation across sex confirmed a significant effect of Drug treatment condition [F (2, 28) = 3.95; P<0.05], but not of Sex or of the interaction of factors. The post-hoc test of the marginal means confirmed that latency was increased after the first THC exposure compared with the PG exposure.

An analysis of the rectal temperature 30 minutes after initiation of inhalation across sex confirmed significant effects of Sex [F (1, 14) = 11.68; P<0.005], of Drug treatment condition [F (2, 28) = 122.5; P<0.0001] and of the interaction of factors [F (2, 28) = 24.01; P<0.0001]. The post-hoc test confirmed that body temperature differed across sexes for all three sessions. The post-hoc also confirmed that female rat temperature in all three conditions differed significantly from each other. Male rat temperature was significantly lower than in the PG condition after each THC condition, which did not differ from each other.

### 3.2 Experiment 2 (Thrice-daily THC inhalation)

THC inhalation decreased body temperature in both male and female groups (Figure 2). The analysis confirmed significant effects of Sex [F (1, 14) = 10.16; P<0.01], Drug treatment condition [F (3, 42) = 394.6; P<0.0001] and the interaction of factors [F (3, 42) = 13.53; P<0.0001]. The post-hoc test confimed that temperature was significantly lower than baseline or PG temperature after each THC condition in both groups. The post-hoc test further confirmed there was a significant attenuation of the hypothermia after THC session 10 versus session 1 for each group and also a sex difference after the first and tenth THC inhalation sessions.

**Figure 2:**
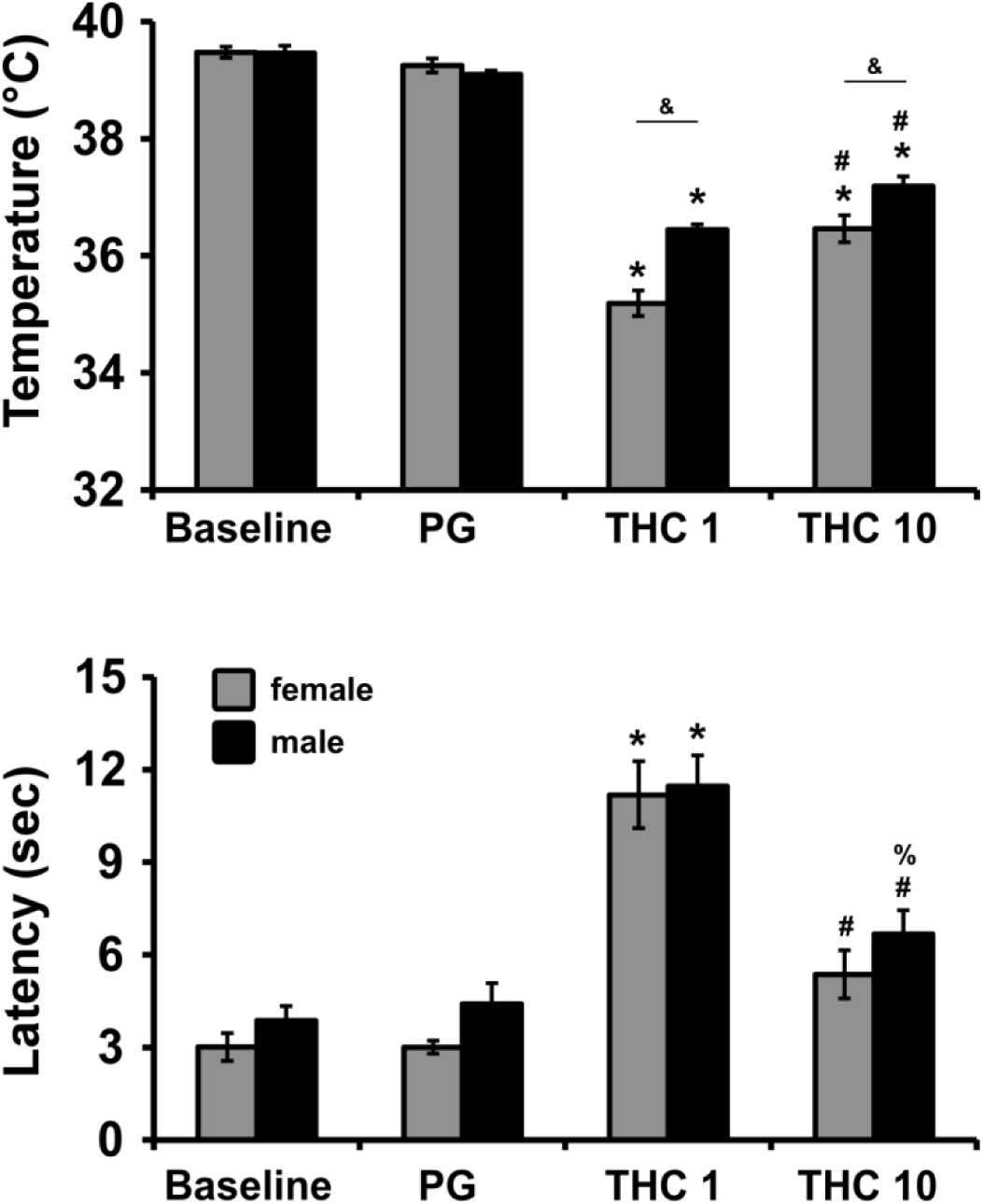
Mean (N=8 per group; ±SEM) rectal temperature (upper panels) and tail-withdrawal latency (lower panels) for male and female rats under baseline conditions or after inhalation of the PG vehicle and then THC (200 mg/mL) on the 1^st^ or 10^th^ repeated session. Significant differences from the Baseline and PG condition are indicated with *, from the Baseline (only) by %, and from the first THC Session with #. A difference between the sexes is indicated with &.

THC inhalation increased antinociception in both male and female groups (Figure 2) but there was a significant attenuation of this effect after repeated THC exposure. The analysis confirmed significant effect of Drug treatment condition [F (3, 42) = 62.4; P<0.0001], but not of sex or the interaction of factors. The post-hoc test confirmed that tail-withdrawal latency was higher after the first THC inhalation compared to all other treatment conditions in each group.

### 3.3 Experiment 3 (Pharmacokinetics)

The mean plasma levels achieved five minutes after the end of a 30 minute session of inhalation were similar for female (303 ng/mL; SEM 37.4) and male (362 ng/mL; SEM 52.5) rats with one significant outlier male contributing to the mean difference (Figure 3). The unpaired t-test did not confirm any differences between the groups. There were no consistent differences associated with the timing of the blood sampling experiment relative to the chronic exposure days (two-way ANOVA not significant). Thus, if there was residual THC in the animals, these levels were negligilble compared to the magnitude of the effect of the acute dosing.

**Figure 3:**
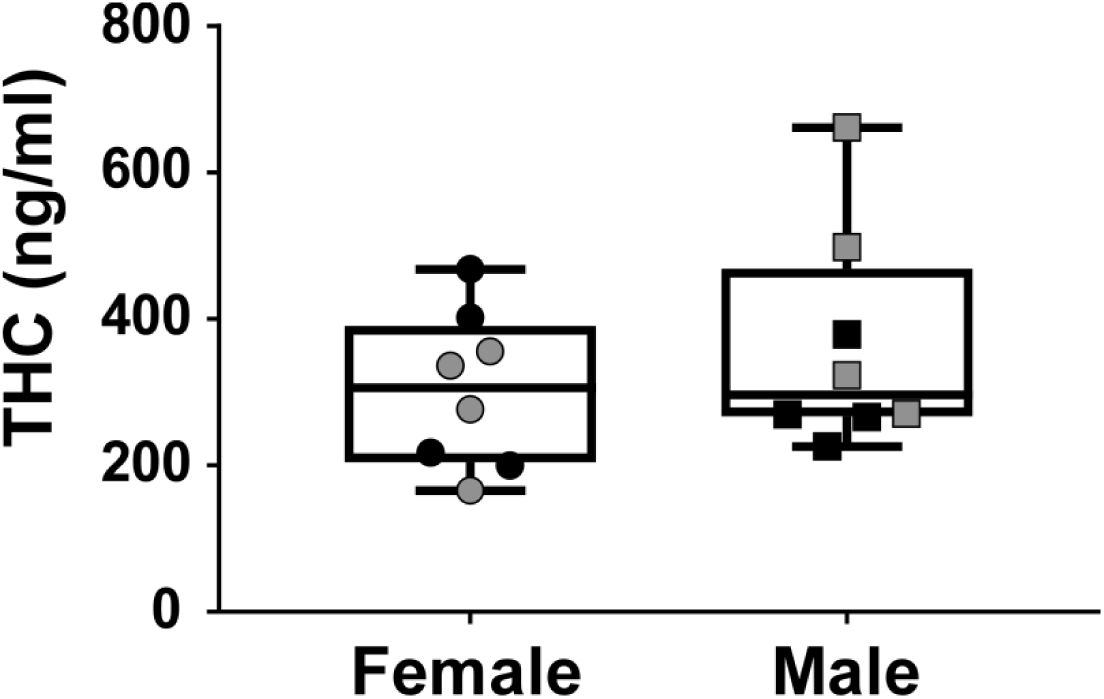
Mean (N=8 per group; ±SEM) plasma THC levels for male and female rats 35 minutes after the start of inhalation of THC (200 mg/mL). Grey symbols indicate the individuals assessed one week after the chronic regimen. Box plot depicting the median and quartiles; the bars indicate the range.

## 4. Discussion

This study is the first to show that repeated inhalation of THC vapor, using an e-cigarette based delivery system, induces tolerance to hypothermia and antinociception in both male and female rats. Significant tolerance was observed following the 7^th^ session of a twice daily regimen in female rats only in the first experiment, and after the 10^th^ session of the t.i.d. regimen in male and female rats in the second experiment. These behavioral results were produced by an acute exposure that produces similar plasma THC leves across male and female animals, as evidenced by Figure 3 and our prior publication (Javadi-Paydar et al., 2017), thus the difference was not attributable to a difference in the individual per-exposure dose. These studies thus demonstrate conditions under which tolerance to the thermoregulatory and antinociceptive effects of vapor inhalation of THC can be produced, further validating the model for investigations of the effects of inhaled THC exposure.

The development of tolerance to THC depends on dose, frequency of exposure and the behavioral or physiological endpoint that is under investigation. One interesting outcome of the study is that tolerance to antinociceptive and thermoregulatory effects of THC inhalation are not necessarily the same in magnitude. In both experiments the relative effect on antinociceptive responses was greater. Thus the primary importance of these data are the demonstration that tolerance can be produced. It has been previously demonstrated that THC exposure by injection or by inhalation approximately once per week does not cause significant tolerance in rats (Borgen and Davis, 1973; Javadi-Paydar et al., 2017). A number of publications show, alternately, that a four to ten day regimen of once or twice daily injection of THC (Delatte et al., 2002; Kwilasz and Negus, 2012; Marusich et al., 2015; Moore et al., 2010; Wakley et al., 2014; Wiley and Burston, 2014; Winsauer et al., 2015), or CB1 full agonists (De Vry et al., 2004), produces tolerance on a variety of measures affected by acute THC. Likewise, tolerance to the effects of inhalation of marijuana smoke was observed after the 13^th^ of every other day exposure (Fried, 1976) in rats but once daily marijuana smoke exposure for two weeks (M-F) did not induce cross tolerance to the locomotor suppressive effects of the endogenous cannabinoid anandamide (Bruijnzeel et al., 2016).

Prior work has shown that female mice develop tolerance to locomotor stimulant effects of THC (i.p.) under conditions under which males do not (Wiley, 2003) and that female rats develop a greater degree of nociceptive tolerance to THC even when the repeated dose is only 71% as large as the male dose (Wakley et al., 2014). Thus an interpretation of the outcome of the present study that male rats may require a more intensive exposure regimen (or a higher dose) to develop tolerance is consistent with prior findings. Despite the piloting of conditions to produce similar effects on temperature and nociception with the second generation apparatus in Experiment 2, the range of blood levels of THC from 303 ng/mL in the female group to 362 (319 with one outlier removed) ng/mL in the male group was higher than the 137 ng/mL found by Javadi-Paydar et al. (Javadi-Paydar et al., 2017) using the first generation equipment and 30 min of inhalation of THC (200 mg/mL). In this study, the magnitude of hypothermia and antinociception on the first exposure to THC was slightly greater in Experiment 2 than in Experiment 1 and this may therefore confirm a difference in plasma THC achieved using the two sets of apparatus. Nevertheless, the magnitude of tolerance in the female rats was, if anything, slightly lesser in Experiment 2 which may reflect the higher probe dose rather than a difference in the development of tolerance *per se*.

Overall this work further confirms the efficacy of a new electronic-cigarette based method of delivering THC to rats. Since humans are increasingly using e-cigarettes to use cannabis extracts, it is necessary to develop pre-clinical models for evaluation of the effects of THC (and other cannabinoids) with this route of administration. This study confirms that tolerance to the hypothermic and antinociceptive effects of THC can be produced with this method. Thus, this method offers excellent face and construct validity for the investigation of the consequences of vapor inhaled THC.

## Author Contributions

MAT and JDN designed the studies, with refinements contributed by TK and YG. MC created the vapor inhalation equipment that was used. JDN, YG, TK and AG performed the research and conducted initial data analysis. JDN and MAT conducted statistical analysis of data, created figures and wrote the paper. All authors approved of the submitted version of the manuscript.

## Acknowledgements

This work was supported by USPHS grants R01 DA035482 (Taffe, PI), R44 DA041967 (Cole, PI) and R01 DA042595 (Cheer, PI). The National Institutes of Health / NIDA had no direct influence on the design, conduct, analysis or decision to publication of the findings. LJARI likewise did not influence the study designs, the data analysis or the decision to publish findings. The authors are grateful to Shawn M. Aarde for significant contributions to the invention and initial validation of the vapor inhalation method and to Mr. Howard Britton for prototyping inhalation equipment. This is manuscript #29555 from The Scripps Research Institute.

